# Multimodal phenotypic axes of Parkinson’s disease

**DOI:** 10.1101/2020.03.05.979526

**Authors:** Ross D. Markello, Golia Shafiei, Christina Tremblay, Ronald B. Postuma, Alain Dagher, Bratislav Miŝić

**Affiliations:** McConnell Brain Imaging Centre, Montréal Neurological Institute, McGill University, Montréal, Canada

**Keywords:** Parkinson’s disease, multimodal fusion, biotype, phenotypic axis, neuroimaging

## Abstract

Individuals with Parkinson’s disease present with a complex clinical phenotype, encompassing sleep, motor, cognitive, and affective disturbances. However, characterizations of PD are typically made for the “average” patient, ignoring patient heterogeneity and obscuring important individual differences. Modern large-scale data sharing efforts provide a unique opportunity to precisely investigate individual patient characteristics, but there exists no analytic framework for comprehensively integrating data modalities. Here we apply an unsupervised learning method—similarity network fusion—to objectively integrate MRI morphometry, dopamine active transporter binding, protein assays, and clinical measurements from *n* = 186 individuals with *de novo* Parkinson’s disease from the Parkinson’s Progression Markers Initiative. We show that multimodal fusion captures inter-dependencies among data modalities that would otherwise be overlooked by field standard techniques like data concatenation. We then examine how patient subgroups derived from fused data map onto clinical phenotypes, and how neuroimaging data is critical to this delineation. Finally, we identify a compact set of phenotypic axes that span the patient population, demonstrating that this continuous, low-dimensional projection of individual patients presents a more parsimonious representation of heterogeneity in the sample compared to discrete biotypes. Altogether, these findings showcase the potential of similarity network fusion for combining multimodal data in heterogeneous patient populations.

## INTRODUCTION

Individuals with Parkinson’s disease (PD) present with a range of symptoms, including sleep, motor, cognitive, and affective disturbances [68]. This heterogeneity is further complicated by individual differences in the age of disease onset and the rate of pathological progression [38, 84]. Most attempts to resolve heterogeneity in PD rely on clustering or subtyping of patients based solely on clinical-behavioral assessments. While these efforts have shown that it is possible to stratify patients into clinically-meaningful categories with considerable predictive utility [22, 24; cf., 74], clinical measures do not directly measure the underlying pathophysiology of PD [21]. Given the prominence of synucleinopathy [29, 50], dopamine neuron dysfunction [23], and distributed grey matter atrophy in PD [93, 96, 97], it is increasingly necessary to develop biologically-informed biotypes by integrating multiple sources of evidence in addition to clinical assessments.

Modern technological advances and data-sharing efforts increasingly permit deep phenotyping in large samples of patients, making simultaneous behavioral assessments, physiological measurements, genetic assays, and brain imaging available at unprecedented scales [53]. A principal challenge is to parsimoniously integrate these multi-view data in order to take full advantage of each source of information [57]. How to account for multiple sources of data to characterize heterogeneous patient samples is a topic of significant interest in computational medicine [40, 58], with important applications for oncology [62, 87], psychiatry, [35, 55, 76], and neurology [21]. Indeed, recent advances in techniques like multiple kernel learning have yielded promising results for integrating disparate data modalities in the context of supervised and unsupervised problems [19, 99]. More broadly, there exist many families of techniques for investigating multi-view data, including multiple kernel learning, matrix factorization, and deep learning, that have been increasingly used in recent years to tackle issues of data integration [100]. Yet, the most commonly employed technique—to simply concatenate data modalities—ignores the structure inherent in individual modalities, potentially yielding biased estimates [87, 100].

In the present report we seek to generate a comprehensive, multi-modal characterization of PD. We apply an unsupervised learning technique, similarity network fusion (SNF; [87]), to integrate data from four data modalities in *n* = 186 individuals with *de novo* Parkinson’s disease from the Parkinson’s Progression Markers Initiative database [53]. Using structural T1-weighted magnetic resonance imaging (MRI), clinical-behavioral assessments, cerebrospinal fluid assays, and single-photon emission computed tomography (SPECT) data we generate patient similarity networks and combine them via an iterative, non-linear fusion process. We demonstrate that SNF yields a more balanced representation of multimodal patient data than standard techniques like data concatenation. We use the patient network generated by SNF to reveal putative PD biotypes, and examine how neuroimaging data contributes to cluster definition and patient discriminability. Finally, we explore how a continuous low-dimensional representation of the fused patient network yields individual estimates of PD patient pathology.

## RESULTS

Complete data were obtained for *n* = 186 patients from the Parkinson’s Progression Markers Initiative database [53]. Data sources included (1) cortical thickness, (2) subcortical volume, (3) clinical-behavioral assessments, (4) dopamine activate transporter (DAT) binding scans, and (5) cerebrospinal fluid assays. Although cortical thickness and subcortical volume are both derived from the same data modality (i.e., anatomical, T1-weighted MRI scans), they are estimated using different algorithms so we retain them as separate sources. For more detailed information on data collection and estimation of derivatives please see *Materials and Methods*.

### Analytic overview

We combine multimodal data using SNF, which first constructs patient similarity networks for each modality and then iteratively fuses the networks together (Fig. 1a). Both the construction of patient similarity networks and the fusion process of SNF are governed by two free hyperparameters. The first parameter, *K*, controls the size of the patient neighborhoods to consider when generating the similarity networks: smaller values of *K* will result in more sparsely connected networks, while larger values will generate denser networks. The second parameter, *μ*, determines the weighting of edges between patients in the similarity network: small values of *μ* will keep only the strongest edges in the patient networks, while larger values will retain a wider distribution of edge weights. We use the fused networks to generate categorical representations of patient data via spectral clustering and continuous representations via diffusion map embedding (Fig. 1b-d). The remainder of the reported results examine the utility of these representations in estimating patient pathology and disease severity.

**Figure 1.**
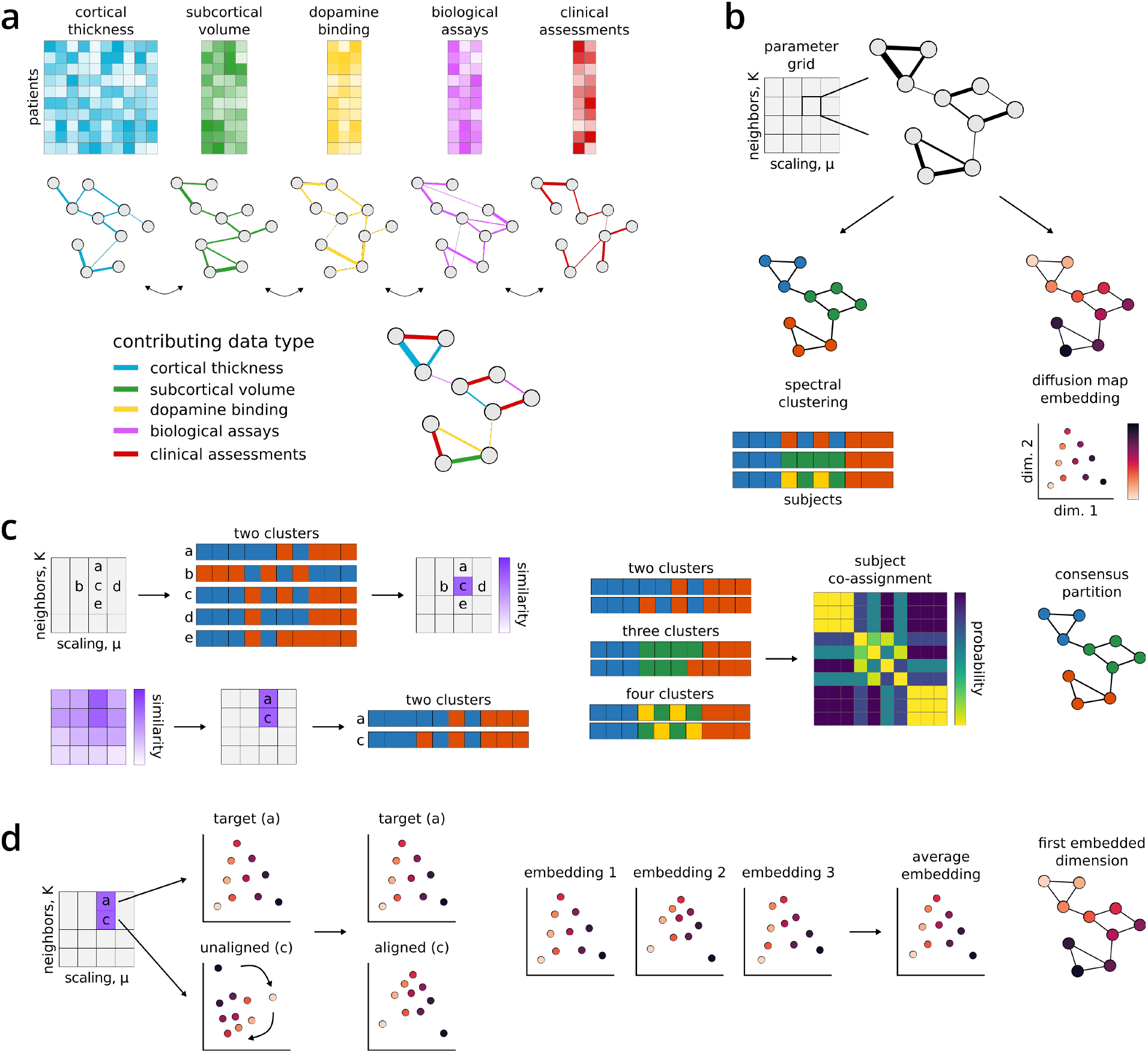
Similarity network fusion analysis pipeline. Toy example demonstrating the processing steps employed in the reported analyses; refer to *Materials and Methods: Similarity network fusion* for more detailed information. Patients are represented as nodes (circles) and the similarity between their disease phenotype is expressed as connecting edges. **(a)** Similarity network fusion generates patient similarity networks independently for each data type and then iteratively fuses these networks together. The resulting network represents patient information and relationships balanced across all input data types. **(b)** We perform an exhaustive parameter search for 10,000 combinations of SNF’s two hyperparameters (*K* and *μ*). The resulting fused patient networks are subjected to (1) spectral clustering and (2) diffusion map embedding to derive categorical and continuous representations of patient data, respectively. **(c)** We assess the local similarity of patient cluster assignments in parameter space using the z-Rand index [80]. The z-Rand index is calculated for all pairs of cluster solutions neighboring a given parameter combination [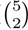 = 10] and then averaged to generate a single “cluster similarity” metric. Clustering solutions from regions of parameter space with an average cluster similarity exceeding the 95th percentile are retained and combined via a consensus analysis to generate final patient clusters (see *Materials and Methods: Consensus clustering*) [6, 46]. **(d)** Diffusion map embedding yields phenotypic “dimensions” of patient pathology [13]. Embeddings from stable regions of parameter space chosen in (c) are aligned via rotations and reflections using a generalized Procrustes analysis and averaged to generate a final set of disease dimensions.

#### Similarity network fusion provides a viable alternative to data concatenation

As the current field standard, data concatenation seems an intuitively appealing method for examining multimodal patient data. In a concatenation framework, features from all data modalities are joined together to create a single patient by feature matrix which is then converted into a patient similarity network using an affinity kernel (e.g., cosine similarity, a radial basis function). This approach is particularly convenient in that it is almost entirely data-driven, with no free parameters beyond the choice of affinity kernel. However, data concatenation tends to suffer from the curse of dimensionality [8]. That is, modalities with many features— regardless of their relative importance—tend to dominate the resulting patient network, obscuring information in modalities with fewer features (Fig. 2a). This is of particular concern when integrating metrics derived from MRI images, which often have tens or hundreds of time more features than lower-dimensionality data sources.

**Figure 2.**
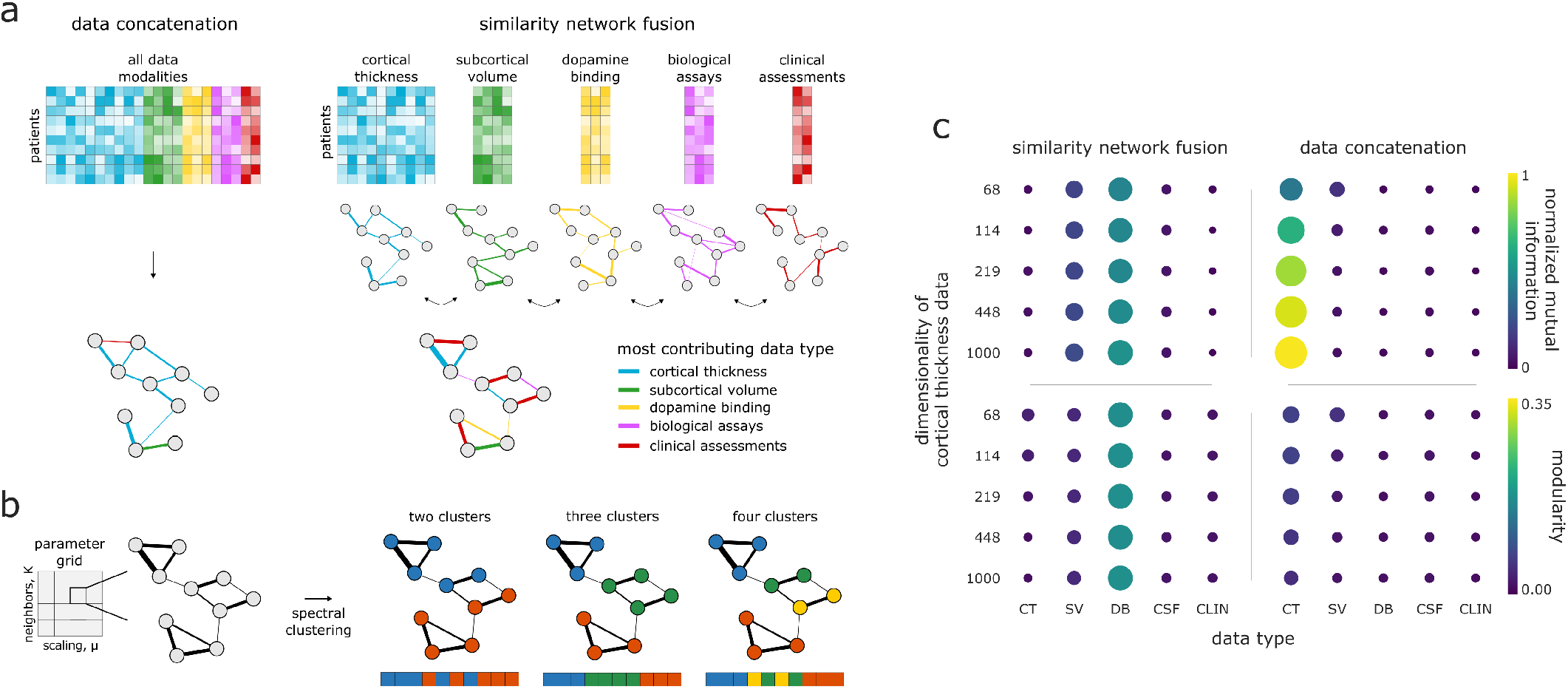
Comparison of data concatenation and SNF for integrating data. **(a)** Toy diagram depicting generation of a patient similarity network from data concatenation in contrast to SNF. The highest dimensionality data—in this case, cortical thickness— tends to be over-represented in the similarity patterns of the patient network generated with data concatenation, whereas SNF yields a more balanced representation. **(b)** An exhaustive parameter search was performed for 10,000 combinations of SNF’s two hyperparameters (*K* and *μ*). We subjected the resulting fused patient network for each combination to spectral clustering for a two-, three-, and four-cluster solution. Clustering solutions were combined via a “consensus” clustering approach (see *Materials and Methods: Consensus clustering*, Fig. 1c) [6, 46]. **(c)** The impact of cortical thickness feature dimensionality on normalized mutual information (NMI) and modularity for different data modalities in data concatenation compared to SNF. NMI estimates are computed by comparing the clustering solutions generated from the concatenated/SNF-derived patient network with the solutions from each single-modality patient network. Modularity estimates are computed by applying the clustering solutions from the concatenated/SNF-derived patient network to the single-modality patient network. CT = cortical thickness; SV = subcortical volume; DB = DAT binding; CSF = CSF assays; CLIN = clinical-behavioral assessments

Similarity network fusion, on the other hand, does not necessarily suffer from such issues. In an SNF framework, patient similarity networks are generated separately for each data modality and then iteratively fused together to create a single, multimodal patient network (Fig. 2a). Converting to patient networks prior to fusing across sources reduces the likelihood of biasing results towards higher-dimensionality data, potentially yielding more balanced representations of the input data [87]. However, SNF is governed by two free hyperparameters, *K* and *μ*, demanding greater computational complexity to avoid arbitrary selection.

To investigate the extent to which SNF is a plausible alternative to data concatenation we generated patient networks using both techniques, varying the dimensionality of cortical thickness data across five increasingly high-resolution subdivisons of the Desikan-Killiany parcellation (ranging from 68–1000 features; see *Materials and Methods: Cortical thickness*) [18].

Rather than selecting a single combination of parameters to combine data modalities in SNF we conducted an exhaustive parameter search, performing the fusion for 10,000 combinations of *K* and *μ* (100 values for each parameter; Fig. 2b). We subjected the resulting networks to spectral clustering with a two-, three-, and four-cluster solution [73, 94]. We then integrated the resulting 30,000 clustering assignments using an adapted consensus clustering approach, previously described in [6] and [46] (see Fig. 1c or *Materials and Methods: Consensus clustering* for more details). Briefly, we assessed the stability of the clustering solutions in parameter space, retaining only those solutions above the 95th percentile of stability, and used the resulting solutions (*n* = 1,262) to create a co-assignment probability matrix which was thresholded and partitioned to generate a set of “consensus” assignments [6, 9].

To assess the relative contribution of the individual data modalities to the concatenation- and SNF-derived clustering assignments we also clustered unimodal patient networks, created separately for each data source (*n* = 5). We compared the similarity between uni- and multimodal clustering assignments using normalized mutual information scores (NMI; [77]), and the goodness-of-fit of clustering assignments to patient networks using modularity [61]. To ensure that the results of the two data integration techniques were more directly comparable, we only used the clustering assignments from concatenated data that had the same number of clusters as those derived via the consensus SNF approach.

We find that at even the lowest dimensionality (i.e., 68 features) patient networks generated using concatenation are dominated by information from cortical thickness data (Fig. 2b). That is, NMI scores show high overlap between clustering assignments derived from only cortical thickness data and assignments generated using concatenated data (average NMI = 0.77 ± 0.25 [0.40–1.00]). Indeed, at the highest cortical thickness dimensionality (1000 features) the clustering assignments for the cortical thickness and concatenated data are identical, suggesting the concatenated patient networks are discarding information from all other lower-dimensionaldata sources. Estimates of modularity are slightly higher for patient networks derived from only cortical thickness data than from other datatypes, but remain low for all data modalities (average modularity = 0.02 ± 0.02 [0.00–0.06]).

On the other hand, SNF appears much more stable to changes in data dimensionality, with NMI scores and modularity estimates more evenly distributed across data modalities (NMI = 0.15 ± 0.18 [0.01–0.50]; modularity = 0.05 ± 0.06 [0.00–0.17]). While this does not necessarily imply that the generated clustering assignments are meaningful, it does suggest that SNF provides a more balanced representation of the input data than simple concatenation, opening the door for a more holistic assessment of their clinical relevance to patient pathology. We chose to examine the fused patient networks generated from the highest (i.e., 1000 node) resolution cortical thickness data for further analyses.

#### Derived patient biotypes are clinically discriminable across modalities

Consensus clustering of the multimodal SNF-derived patient networks yielded three clusters—or “biotypes”— of *n* = 72, 69, and 45 individuals (Fig. 3a,b,c) with strong goodness-of-fit (modularity = 0.494, *p* < 0.001 by permutation). There were no significant inter-group differences for sex (*p* = 0.32), age (*p* = 0.30), education (*p* = 0.77), symptom duration (*p* = 0.83), recruitment site (*p* = 0.20), or MRI scanner strength (*p* = 0.38) (see Table S2 for summary demographics of each cluster).

**Figure 3.**
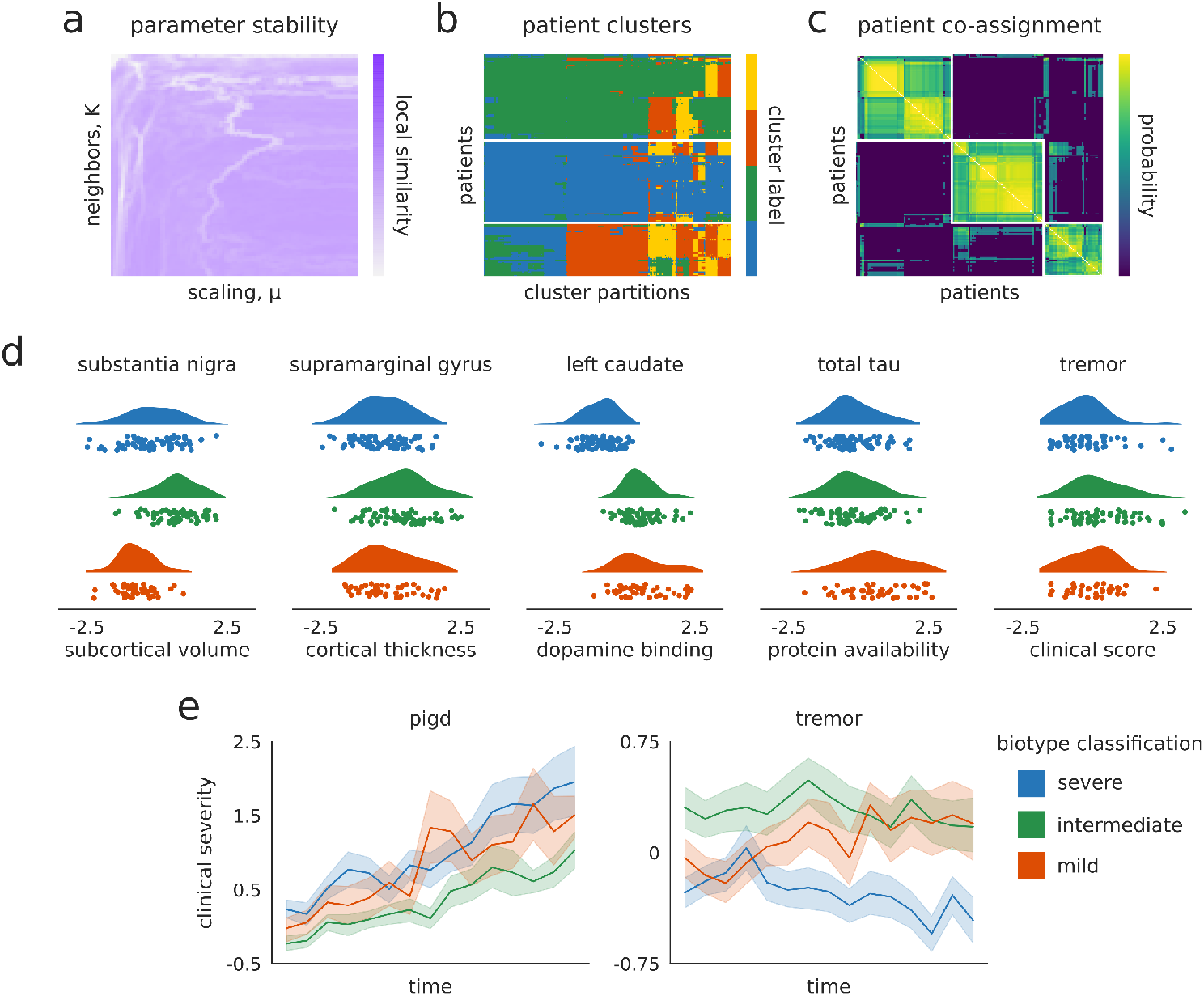
Patient biotypes discriminate multimodal disease severity. **(a)** Local similarity of clustering solutions for examined SNF parameter space, shown here for the three-cluster solution only. **(b)** Patient assignments for all of the two-, three-, and four-cluster solutions extracted from stable regions of the SNF parameter space in (a). The three cluster solution is ostensibly the most consistent across different partitions. **(c)** Patient co-assignment probability matrix, indicating the likelihood of two patients being assigned to the same cluster across all assignments in (b). The probability matrix is thresholded based on a permutation-based null model and clustered using an iterative Louvain algorithm to generate consensus assignments (see *Materials and methods: Consensus clustering*). **(d)** Patient biotype differences for the most discriminating feature of each data modalities at baseline. X-axis scales for all plots represent z-scores relative to the sample mean and standard deviation. Note that all ANOVAs with the exception of tremor scores are significant after FDR-correction (q < 0.05). Higher cortical thickness, subcortical volume, and DAT binding scores are generally indicative of clinical phenotype; refer to Fig. S1 for guidelines on how biological assays and clinical-behavioral assessments coincide with PD phenotype. **(e)** Patient biotype differences for two relevant clinical-behavioral assessments over time; shaded bands in (e) indicate standard error. Linear mixed effect models were used to examine the differential impact of biotype designation on longitudinal progression of clinical scores; refer to Table S4 for model estimates.

To assess the clinical relevance of these biotypes we performed a series of univariate one-way ANOVAs for all 1,050 input features (see Table S1 for a complete list). We found 31 features that significantly distinguished patient groups from one another (false discovery rate [FDR] corrected, q < 0.05; the most discriminating feature from each data modality are shown in Fig. 3d and a full list is available in Table S3). Though biotypes showed limited differences in clinical-behavioral assessments using baseline data, longitudinal analyses of subgroup affiliation revealed differentiation over time in clinical measurements frequently used for PD prognosis (Fig. 3e; refer to Table S4 for model estimates). Note that while increases in both tremor and postural instability/gait difficulty (PIGD) scores are indicative of clinical severity, higher PIGD (and lower tremor) scores are often found to be related to a more rapidly progressing and severe manifestation of PD [36].

Broad examination of the DAT binding, CSF, and clinical assessment data suggests that the three biotypes may separate along a single dimension of PD severity, where group one represents a more severe, group two an intermediate, and group three a more mild phenotype; however, differences in the neuroimaging data reveal this delineation is less straightforward. For instance, individuals in group two—the ostensible “intermediate” clinical biotype—also tend to have greater subcortical volume than individuals in the “mild” clinical group, especially in brain regions typically prone to degeneration in early PD such as the substantia nigra (*F*(2, 183) = 40.22, *p* = 8.70 x 10^−13^; Fig. 3d). That is, assignment of cluster labels indicative of disease severity are incapable of capturing all phenotypic aspects of PD.

To further investigate the discrepancy between the clinical and neuroimaging data, we calculated patient scores for a putative neuroimaging biomarker—the “PDICA atrophy network”—recently shown to have relevance to both diagnostic PD disease severity and longitudinal prognosis [95] (Fig. 4c). While the PD-ICA biomarker was generated from the same dataset examined in the current study, it was derived via metrics which we excluded from our SNF analyses (i.e., whole-brain deformation-based morphometry [DBM] estimates). In line with results from the neuroimaging data included in SNF, we found significant differentiation of PD-ICA atrophy scores between biotypes (*F*(2, 183) = 5.70, *p* = 0.004), largely driven by lower atrophy in the “intermediate” compared to the “mild” group (post-hoc Tukey test, p < 0.05). These observed discrepancies between MRI-derived metrics and other modalities raise the question: how important is MRI brain imaging to characterizing PD pathology?

**Figure 4.**
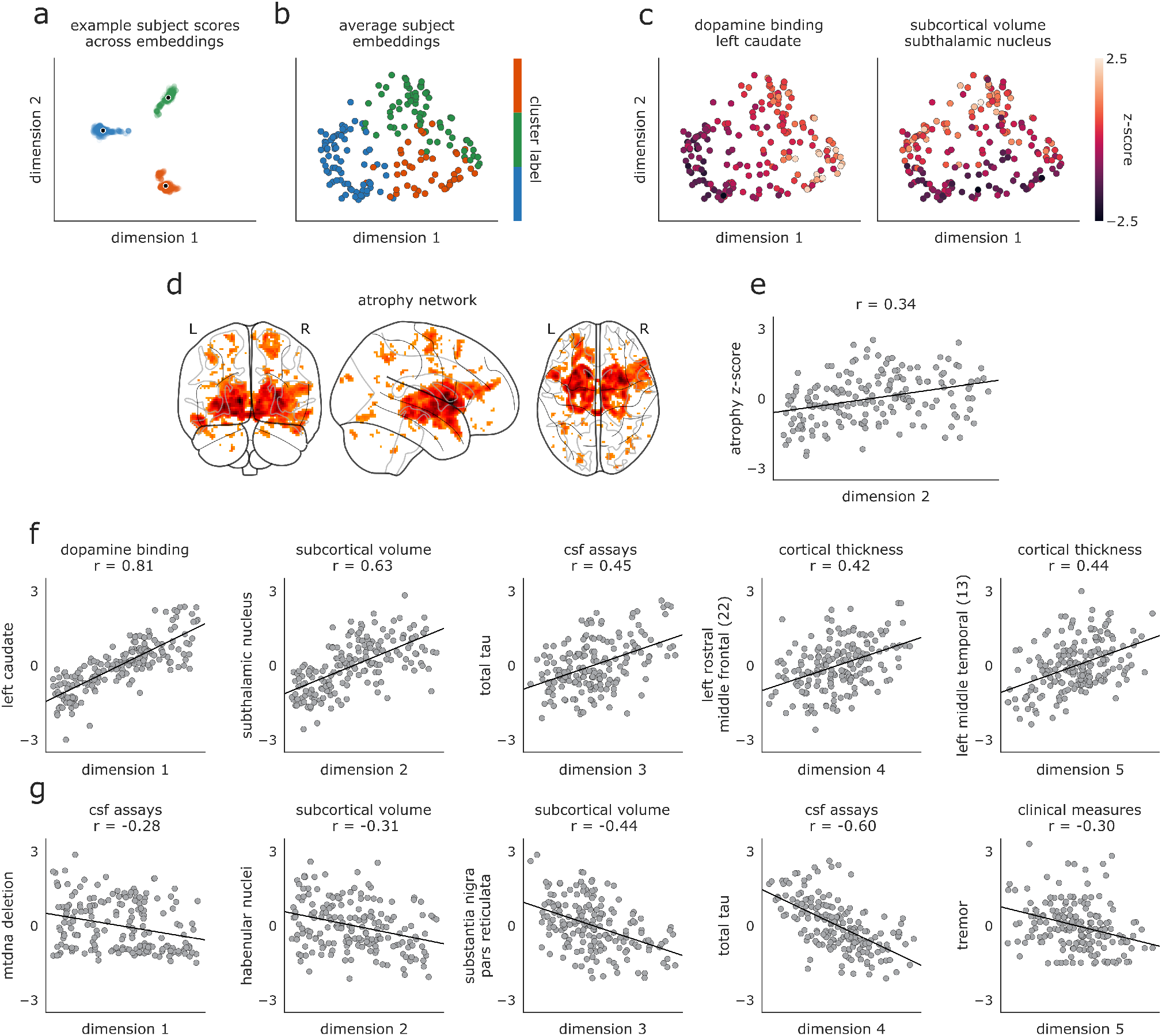
Embedded patient dimensions capture relevant clinical information. **(a)** Scores of three example patients (one from each biotype) along the first two embedding dimensions generated from all 1,262 hyperparameter combinations retained in analyses; black dots represent average scores across all parameter combinations, which are shown for every patient in (b). **(b)** Scores of all patients along first two dimensions of average embeddings. Colors reflect same biotype affiliation as shown in Fig. 3. **(c)** Same as (b) but colored by patients scores of the feature most strongly correlated with the first and second embedding dimensions. Left: colors represent DAT binding scores from the left caudate (*r* = 0.81 with dimension one). Right panel: colors represent subcortical volume of the subthalamic nucleus (*r* = 0.63 with dimension two). **(d)** PD-ICA atrophy network from [97], showing brain regions with greater tissue loss in PD patients than in age-matched healthy individuals. **(e)** PD atrophy scores, calculated for each patient using the mask shown in (e), correlated with subject scores along the second embedded dimension shown in (a-c). **(f, g)** Features with the most positive (f) and negative (g) associations to the five embedding dimensions. Y-axes are presented as z-scores; x-axes are dimensionless units derived from the diffusion embedding procedure.

#### Neuroimaging data is critical to patient biotype characterization

Although one benefit of SNF is the ability to seamlessly integrate neuroimaging data with clinical assessments, high-resolution MRI scans can be both costly and inconvenient for PD patients. Most previous biotyping and patient classification studies in PD have focused solely on clinical-behavioral assessments [22, 24], raising the possibility that an adequate solution can be identified without MRI scans. Thus, we investigated whether a similar patient characterization can be identified using a reduced dataset excluding MRI data.

Data from clinical-behavioral assessments, cerebrospinal fluid assays, and subcortical DAT binding scans were used to generate fused patient networks following the same procedures described above (see *Similarity network fusion provides a viable alternative to data concatenation*). Consensus clustering of the networks yielded three subgroups of *n* = 83, 59, and 44 individuals, showing moderate overlap with biotypes defined on the full dataset (NMI = 0.43).

Despite this similarity, it is possible these “no-MRI” biotypes would result in a different characterization of PD disease severity than the previously described biotypes. To assess this, we applied the same univariate one-way ANOVA framework to examine which of the original 1,050 features were discriminable between the subgroups generated without MRI data. This revealed only five features (from two modalities) that significantly distinguished these patient subgroups (FDR-corrected, q < 0.05): DAT binding in bilateral caudate and putamen and clinical scores on the the Unified Parkinson’s disease rating scale, part II (UPDRS-II; [30]). Comparing these features with the complementary set derived from all the data (Table S3) highlights the reduced discriminability of these “no-MRI” subgroups.

Given that the reduced dataset contains relatively few (i.e., 34) features, it is possible that SNF may be “overengineering” a solution that could be better achieved with alternative clustering techniques. To address this possbility, we reproduce a previously published clustering solution of PD patients that does not include features derived from MRI data and compare the results of this technique to SNF (see *Supplementary Materials and Results: Comparing clustering techniques*). We find that, as above, this clustering solution fails to yield subgroups with meaningful differences for any neuroimaging metrics.

Taken together, these results support the notion that excluding MRI data from PD clustering yields patient subgroups that fail to capture significant differences in PD pathophysiology. Nevertheless, the value of neuroimaging data in defining patient subgroups does not explain the inconsistencies previously observed between neuroimaging metrics and clinical data in the patient biotypes. In the early stages of PD, pathophysiology may not perfectly align with clinical symptomatology; that is, hard partitioning into a small number of discrete biotypes may be an over-simplification of the disease.

#### Patient biotypes separate along continuous dimensions of severity

To examine the possibility that PD pathology is multi-dimensional, we generated a continuous lowdimensional representation of our fused patient networks with diffusion embedding [13, 44]. Diffusion embedding attempts to find dimensions (often referred to as gradients or components) of a network that encode the dominant differences in patient similarity—akin to principal components analysis (PCA) or multidimensional scaling (MDS). The resulting components are unitless and represent the primary axes of inter-subject similarity. Critically, diffusion embedding is sensitive to complex, non-linear relationships and is relatively robust to noise perturbations compared to other dimensionality reduction methods [13, 44].

Patient networks for all SNF hyperparameter combinations used in the consensus clustering (*n* = 1,262 networks) were decomposed with diffusion embedding and realigned using a generalized orthogonal Procrustes analysis. The resulting aligned embeddings were then averaged to generate a single, embedded space (Fig. 1d). Examining the extent to which patient clusters differentiated in embedded space revealed limited overlap in cluster affiliation among the first two dimensions (Fig. 4a,b).

The first two dimensions of the embedded space correlated most strongly with patient variability in DAT binding of the left caudate (dimension one: *r* = 0.81; Fig. 4c, left panel) and volume of the subthalamic nucleus (dimension two: *r* = 0.63; Fig. 4c, right panel). Examining further dimensions revealed significant relationships with features from all data modalities (Fig. 4f-g). We also investigated the extent to which patient PD-ICA atrophy scores related to variation along estimated diffusion dimensions [97], finding significant associations between atrophy scores and dimensions two and four (*r* = 0.33 and −0.26, FDR-corrected, *q* < 0.05; Fig. 4d,e).

To examine whether this dimensional framework provided out-of-sample predictive utility we re-ran the SNF grid search excluding two variables frequently used in clinical settings to assess disease severity: the tremor dominant score and the postural instability/gait difficulty score (PIGD; [75]). We regenerated the embedded dimensions following the procedure depicted in Fig. 1d and used five-fold cross validation to assess the extent to which embeddings predicted patient scores on the heldout clinical variables. A simple linear regression was performed using the first five embedding dimensions as predictors of each clinical score; betas were estimated from 80% of the data and applied to predict scores for the remaining 20%. Out-of-sample scores were predicted moderately well (average Pearson correlation between real and predicted scores: *r*_tremor_ = 0.20 [SD = 0.22]; *r*_pigd_ = 0.22 [SD = 0.22]).

## DISCUSSION

The present report demonstrates how diverse modalities can be integrated to comprehensively characterize a range of patient characteristics. Using information about behavior, DAT binding, cerebrospinal fluid, cortical thickness, and subcortical tissue volume, we find evidence for distinct biological dimensions or phenotypic axes that span the patient sample. These biologically-informed dimensions provide a more nuanced interpretation of pathology than is permitted from discrete subgroups or biotypes, and hold potential for greater precision in diagnosis.

### Parkinson’s disease heterogeneity

The present study demonstrates that even in a well-controlled sample of *de novo* PD patients, there exists considerable heterogeneity. This finding contributes to a rich literature on the diversity of PD symptoms [20, 21, 24, 48, 49, 71, 79]. Despite the fact that the putative biotypes are based on the first patient visit, there are several important features where the biotypes are not initially different from each other but progressively diverge over time (e.g., tremor scores).

Does the observed heterogeneity imply the existence of fundamentally different diseases, or simply different rates of progression? Recent evidence from animal models suggests that PD originates from misfolding and trans-synaptic spreading of the endogenous protein *α*synuclein [50], with convergent results from human neuroimaging [28, 64, 93, 98]. The large-scale atrophy patterns and associated neurological deficits are thought to be mediated by the spread of these pathogenic protein aggregates [91], penetrating the cerebral hemispheres via the brainstem and subcortex [89]. The heterogeneity of clinical symptoms may simply suggest that patients may differ in the rate and extent of neurodegeneration, leading to diverse symptoms. In other words, individual variation in neurological manifestations of the disease may depend on the spreading pattern and the affected networks [52], as well as factors not directly related to PD pathophysiology (e.g., age, sex, comorbidities).

### Biotypes or dimensions?

Hard partitioning methods yield non-overlapping clusters by definition, but is this the best way to characterize heterogeneity in PD? The patient biotypes we initially identified could largely be differentiated from each other on the basis of disease severity, leading us to pursue a dimensional approach. We find that much of the phenotypic variability in the patient sample could be parsimoniously captured by a smaller number of latent dimensions or phenotypic axes that relate to distributed patterns of grey matter atrophy, DAT binding, etc. This finding is reminiscent of previous reports, where even hard clustering solutions often sorted patients into broad disease severity categories [20, 24, 48, 49, 79]. Indeed, recent studies have taken an explicitly dimensional approach, attempting to find low-dimensional projections of clinical-behavioral and neuroimaging data [39, 41, 71, 78]. In clinical practice, methods like SNF situate individual patients in a biologically-comprehensive feature space that can then guide more objective clinical decisions about diagnosis and prognosis. Our study is a first step in better understanding which measures are necessary and informative of PD severity, but more work needs to be done to continue to investigate how this will translate to clinical practice.

The notion that PD can be characterized by a smaller number of latent dimensions opens the question of where PD is situated relative to other diseases [5]. Given the natural functional dependencies among the molecular components of a cell, distinct pathological perturbations (e.g. mutations) may affect overlapping gene modules, cell types, tissues and organs, ultimately manifesting in similar phenotypes. For instance, do the present latent dimensions trace out a transdiagnostic continuum along which we can place other neurodegenerative diseases (e.g. Alzheimer’s disease, tauopathies). How PD fits into a global “disease-ome” remains an exciting question for future research.

### Data integration for precision medicine

More broadly, the present work builds on recent efforts to draw insight from multiple modalities. Modern technological advances and data sharing initiatives permit access to large patient samples with increasing detail and depth [53]. How best to integrate information from these multi-view data sets remains a fundamental question in computational medicine [57, 87, 100]. Though there is active discussion regarding what stage of analysis is most appropriate for data integration [100], our results demonstrate that important insights about PD patient heterogeneity can be drawn only by simultaneously considering multiple data modalities. Covariance among clinical, morphometric, and physiological measurements synergistically reveals dominant axes of variance that are not apparent in the individual data modalities. Moreover, we show that simple concatenation induces overfitting to the modality with the greatest dimensionality and autocorrelation, motivating further research on integrating diverse sources of information.

A corollary of the present work is that it is also possible to identify modalities that *do not* make a significant contribution towards differentiating patients and could potentially be excluded from future data collection efforts. This is an important concern because many modern biological assays—such as brain imaging—are expensive and difficult to administer for some types of clinical populations. As a result, samples may be biased and data may be incomplete for many individuals, limiting the application and utility of the subsequent statistical models. The present analysis can thus help to streamline future data collection efforts. By helping to reduce the feature set, the present analytic approach can also help increase potential for overlap and interoperability among existing datasets.

### Methodological considerations

Although we took steps to ensure that the reported results are robust to multiple methodological choices, there are several important limitations to consider. First, the present results are only demonstrated in a single patient sample; despite consistency with several other recent studies [71], formal replication in new longitudinal cohorts is necessary.

Second, the present statistical model leaves out two widely-available and potentially important data modalities: genetic variation and daily movement. Genetic variation is a recognized contributor to the clinical manifestations of PD [60], and previous work shown that genetic risk can be used to meaningfully stratify patients [71]. Likewise, objective measurements of daily movement with wearable sensors and smart devices are increasingly prevalent and add a fundamentally different source of information about individual patients [51]. Whether and to what extent the present phenotypic axes reflect the underlying genetic determinants or movement characteristics of PD is an exciting question for future research.

Finally, there has been recent work highlighting the potential of diffusion-weighted imaging (DWI) as an alternative measure to structural, T1w images for predicting PD prognosis [1]; however, the limited availability of DWI in longitudinal cohorts like those used in the current study restricts its more widespread use.

### Summary

In summary, we report a flexible method to objectively integrate a diverse array of morphometric, molecular and clinical information to characterize heterogeneity in PD. We find evidence for three biotypes, but show that the sample can alternatively be characterized in terms of continuous phenotypic dimensions. These phenotypic dimensions bring into focus complementary information from multiple modalities and lay the foundation for a more comprehensive understanding of Parkinson’s disease.

## MATERIALS AND METHODS

### Code and data availability

Data used in this study were obtained from the Parkinson’s Progression Markers Initiative (PPMI) database (https://www.ppmi-info.org), accessed in March of 2018. All code used for data processing, analysis, and figure generation is available on GitHub (https://github.com/netneurolab/markello_ppmisnf) and relies on the following open-source Python packages: h5py [14], IPython [67], Jupyter [43], Matplotlib [33], NiBabel [10], Nilearn [2], Numba [45], NumPy [63, 85], Pandas [56], scikit-learn [66], SciPy [86], StatsModels [72], Seaborn [90], BCTPy [70], and mapalign [47].

### Neuroimaging data processing

Raw neuroimages were directly converted from DICOM to BIDS format [31] using heudiconv (https://github.com/nipy/heudiconv). Structural images for each subject were then independently processed with the Advanced Normalization Tools’ (ANTs) longitudinal cortical thickness pipeline [83].

Briefly, T1-weighted structural images were corrected for signal intensity non-uniformity with N4 bias correction [81]. Corrected images were then combined with other available neuroimages (T2-weighted, proton-density, and FLAIR images) across all timepoints to create a temporally unbiased, subject-specific template [4]. A standard template was non-linearly registered to the subject template and used to remove nonbrain tissue; to maintain consistency with previous work on the PPMI dataset [93, 97] we used the Montreal Neurological Institute (MNI) ICBM-152 2009c template for this purpose (https://www.bic.mni.mcgill.ca/ServicesAtlases/ICBM152NLin2009; [15, 25, 26]). The remaining subject template brain tissue was segmented into six classes (cerebrospinal fluid, cortical gray matter, white matter, subcortical gray matter, brain stem, and cerebellum) using ANTs joint label fusion [88] with a group of fifteen expertly annotated and labeled atlases from the OASIS dataset [42]. Finally, the segmented subject template was used to performing the same registration, brain extraction, and segmentation procedures on the T1w images from each timepoint.

Three metrics were derived from the pre-processed neuroimaging data: subcortical volume, cortical thickness, and whole-brain deformation-based morphometry values.

#### Subcortical volume

Subcortical volume measures the absolute volume, in cubic millimeters, for pre-defined regions of interest. Here, we used the regions of interest from the highresolution subcortical atlas generated by [65]. Analyses reported in the main text used the deterministic version of the atlas; however, using the probabilistic version returned comparable results. Where applicable the probabilistic atlas was thresholded at 40%; thresholding was performed after registration to subjects’ T1w MRIs.

#### Cortical thickness

Cortical thickness measures the distance, in millimeters, between the pial surface and the gray-white matter boundary of the brain. The ANTs diffeomorphic registration-based cortical thickness estimation (DiReCT) algorithm was used to measure cortical thickness in the volumetric space of each T1w image [16]. The diffeomorphic constraint of this procedure ensures that the white matter topology cannot change during thickness estimations, permitting accurate recovery of cortical depth even in sulcal grooves. Previous work has found that this procedure compares favorably to surface-based approaches [82].

To account for subject-level neuroanatomical variance, we averaged cortical thickness values in each region of the five multi-scale parcellations (68, 114, 219, 448, and 1000 regions) generated by [11] for every timepoint for every subject. The parcellations were transformed to the T1w MRI of each timepoint before averaging within regions to minimize bias.

#### Deformation-based morphometry

Deformation-based morphometry (DBM) provides an approximate measure of local changes in brain tissue volume for a subject relative to a standard template [3]. DBM values are derived from the deformation maps generated during the non-linear registration process aligning each subject brain to the template space. In the present study, deformation maps were created by concatenating the non-linear warps described in *Neuroimaging data processing* that (1) mapped the T1w image for each timepoint to the relevant subject-specific template and (2) mapped the subject-specific template to the MNI152-2009c template. Local changes in tissue volume were then estimated from the derivative of these deformation maps, calculated as the determinant of the Jacobian matrix of displacement. To aid interpretability of these changes we calculated the natural log of the Jacobian determinant, such that a value of zero indicates no volume change compared to the MNI template, negative values indicate tissue expansion relative to the MNI template, and positive values indicate tissue loss relative to the MNI template.

*PD-ICA atrophy calculation* Zeighami and colleagues initially used DBM values from the PPMI subjects to find an ICA component map, which they refer to as a PDICA map, highlighting regions of the brain with greater relative tissue loss in Parkinson’s patients than in age-matched healthy controls [97]. Using this map, they generated an “atrophy score” for each patient, which they related to measures of PD disease severity and prognosis [95, 97]. Unfortunately, a precise calculation of this atrophy score on a different subset of subjects from the PPMI would require the original ICA component table; in order to approximate this score without the component table we employed an alternative procedure originally used in [95].

We downloaded the PD-ICA component map from NeuroVault (collection 860, image ID 12551, https://identifiers.org/neurovault.collection:860; [32]) and used the map as a weighted mask on the DBM images estimated for each subject, multiplying DBM values by the component weights in the map and then summing the resulting values to generate a single score for each time point for each subject. Reported atrophy scores are all normalized (i.e., zero mean and unit variance) with respect to the *n* = 186 patient sample.

#### Quality control

Neuroimaging data were visually inspected by two authors (RDM and CT) using tools adapted from niworkflows (https://github.com/poldracklab/niworkflows). Processed data from twenty individuals were jointly selected by both raters as anchors and used to guide independent rating of the remaining T1w structural images for all subjects [69]. Quality of brain extraction, brain segmentation, and registration to the MNI152-2009c template were assessed and rated on a scale of 0–2, where 0 indicates a processing failure, 1 indicates a conditional pass, and 2 indicates a full pass. Discrepancies where one rater assigned a score of 0 and the other rater assigned a passing score (*n* = 11 instances) were jointly reconciled and new scores assigned (*n* = 9 revised to a score of 0, *n* = 2 revised to a score of 1).

Cohen’s kappa coefficient (*κ*) was calculated to compare the inter-rater reliability of the final quality control scores, yielding 84.8% agreement for segmentation and 84.5% agreement for registration ratings [12]. Assessing scores for only the subject-specific template of each subject yielded comparable agreement (85.8% for segmentation, 85.5% for registration). The current study used subjects for whom both segmentation and registration scores were ≥1 across both raters.

### Non-neuroimaging data processing

In addition to longitudinal neuroimages, the PPMI provides clinical-behavioral assessments, biospecimen analyses, and single-photon emission computed tomography (SPECT) dopamine active transporter (DAT) binding data for all of its participants. Item-level measures for clinical-behavioral assessments were combined into raw composite scores following instructions in the “Derived Variable Definitions and Score Calculations” guide supplied by the PPMI using the pypmi software package (https://github.com/netneurolab/pypmi). Biospecimen data and pre-computed region of interest metrics for SPECT data were used as provided.

### Data cleaning

The current study used five data sources: (1) MRI cortical thickness, (2) MRI subcortical volume, (3) SPECT DAT binding ratios, (4) biological and cerebrospinal fluid assays, and (5) clinical-behavioral assessment scores. Prior to combining data sources we performed (i) data cleaning, (ii) outlier removal, (iii) missing data imputation, (iv-a) batch correction, (iv-b) covariate residualization, and (v) normalization for each modality.

Any features for which ≥20% of individuals were missing data were discarded; subsequently, individuals missing 20% of the remaining features were discarded. Putative outlier individuals were then identified using a median absolute deviation method and discarded [34]. Remaining missing data values were imputed, substituting the median value across individuals for each feature. This procedure yielded *n* = 186 individuals with PD and *n* = 87 healthy individuals who had data from all five sources. A full list of the 1,050 features retained for each data source after preprocessing can be found in Table S1. We used ComBat to correct for site differences in neuroimaging data modalities (i.e., cortical thickness and subcortical volume measurements; neurocombat, https://github.com/ncullen93/neurocombat) [27, 37]. Patient diagnostic status, family history of PD, sex, race, handedness, and education were included in the ComBat correction procedure.

We residualized the pre-processed data against age, sex, and age × sex interactions in PD patients based on the relationships estimated from healthy individuals; estimated total intracranial volume was also included in the residualization process for cortical thickness and subcortical volume features [96]. Finally, PD patient data were z-scored (centered to zero mean and standardized to unit variance). Only PD patient data were used to estimate means and standard deviation for z-scoring.

### Data concatenation

Concatenation provides a data-driven, parameter-free approach for integrating multimodal data, where data features from all sources are horizontally stacked into a single sample by feature matrix. Here, we joined patient data for all features from all data sources and converted the resulting matrix into a patient similarity network by applying a cosine similarity function from scikit-learn [66]. As spectral clustering cannot handle negative values we scaled the values of the similarity networks between zero and two. The scaled patient networks were subjected to spectral clustering [73, 94] for a two-, three-, and four-cluster solution, and the solutions were compared to clustering assignments generated from SNF, described below.

### Similarity network fusion

Similarity network fusion is a method for combining disparate data sources from a group of samples into a single graph, representing the strength of relationships between samples [87]. SNF constructs independent similarity networks from each data source using a K-nearest neighbors weighted kernel and then iteratively combines them via a non-linear message passing protocol. The final network contains sample relationships representing information from all data sources and can be subjected to clustering or other graph techniques (see Fig. 1a). SNF belongs to a broader family of techniques of multi-view learning algorithms (e.g., multiple kernel learning, multi-table matrix factorization). We opted to use SNF because (1) it is an unsupervised learning technique, (2) it is explicitly optimized to control for differing dimensionalities amongst input data modalities [87], and (3) it has been shown to be effective at disentangling heterogeneity in psychiatric populations [35, 76].

#### Network creation and fusion

We constructed similarity networks for each data source using a Python implementation of the methods described in [87] (snfpy; https://github.com/netneurolab/snfpy). A brief description of the main steps in SNF follows, adapted from its original presentation in [87].

First, distance matrices are created from each feature matrix. We selected squared Euclidean distance for the analyses in the main text; however, other distance measures return comparable results (see *Supplementary Materials: Alternative distance metrics*). Next, distance matrices are converted to similarity networks using a scaled exponential kernel:

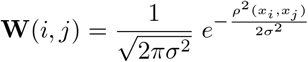

where *ρ*(*x_i_, x_j_*) is the Euclidean distance (or other distance metric, as appropriate) between patients *x_i_* and *x_j_*. The value *σ* is calculated with:

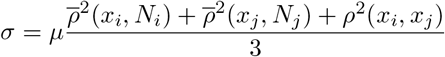

where 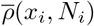 represents the average Euclidean distance between *x_i_* and its neighbors *N*_1*..K*_. Both *K*, controlling the number of neighbors, and *μ*, the scaling factor, are hyperparameters that must be pre-selected, where *K* ∈ [1, 2*, …, j*]*, j* ∈ ℤ and *μ* ∈ ℝ^+^.

In order to fuse the supplied similarity networks each must first be normalized. A traditional normalization performed on a similarity matrix would be unstable due to the self-similarity along the diagonal; thus, a modified normalization is used:

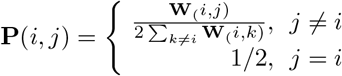

Under the assumption that local similarities are more important or reliable than distant ones, a more sparse weight matrix is calculated:

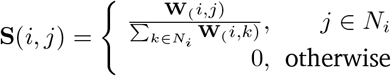

The two weight matrices **P** and **S** thus provide information about a given patient’s similarity to all other patients and the patient’s *K* most similar neighbors, respectively.

The similarity networks are then iteratively fused. At each iteration, the matrices are made more similar to each other via:

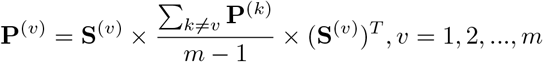

After each iteration, the resulting matrices are renormalized via the above equations. Fusion stops when the matrices have converged or after a pre-specified number of iterations (by default, 20).

#### Hyperparameter search

Variation in SNF’s two parameters, *K* and *μ*, can highlight different aspects of the input data, yielding significantly different fused networks. In order to avoid biasing our results by selecting any specific hyperparameter combination we opted to perform SNF with different combinations of *K* and *μ*, using 100 unique values for each parameter (*K* = 5–105; *μ* = 0.3–10, logspace). The resulting set of 10,000 fused networks was subjected to consensus clustering and diffusion map embedding to generate both categorical and continuous representations of patient pathology.

For a more in-depth examination of how variation in hyperparameters impacted patient networks refer to *Supplementary Materials: Hyperparameter variation*.

### Consensus clustering

In order to find a single clustering solution from the 10,000 fused networks generated by SNF we employed a consensus approach inspired by [6] and [46]. First, each of the 10,000 networks was subjected to spectral clustering [73, 94]. As spectral clustering requires prespecifying the desired number of clusters to be estimated, we chose to generated separate solutions for two, three, and four clusters each. Though it is possible there are more clusters in the dataset, previous work on PD patient data supports 2–4 clusters as a reasonable choice [22, 24].

We wanted to consider regions of the parameter space that generated “stable” networks—that is, where small perturbations in either of the two hyperparameters did not appreciably change the topology or resulting clustering of the patient networks. To quantify this we assessed the pairwise z-Rand similarity index [80] of the clustering solution for each point in hyperparameter space and its four neighbors; local z-Rand values were calculated separately for two-, three-, and four-cluster solutions (Fig. 1c). The resulting cluster similarity matrices were thresholded at the 95th percentile of the values in all three matrices; clustering solutions from regions of parameter space surviving this threshold in any of the three matrices were retained for further analysis, resulting in 1,262 3 = 3,786 assignments. These solutions were used to generate a subject co-assignment matrix representing the normalized probability that two subjects were placed in the same cluster across all assignments. This “co-assignment” matrix was thresholded by generating an average probability from a permutation-based null model, as in [6]. Briefly, we permuted the assignments for each clustering solution and re-generated the co-assignment probability matrix; the average probability of this permuted matrix was used as the threshold for the original. We clustered the resulting thresholded matrix using a modularity maximization procedure to generate a final “consensus” clustering partition, which was used in all subgroup analyses [9].

### Diffusion map embedding

While clustering is appealing for its intuitive clinical applications, recent work has shown that continuous representations of clinical dysfunction may provide a more accurate representation of the underlying diseases [92, 96] As an alternative to clustering, we applied diffusion embedding to the fused PD patient network.

Diffusion map embedding is a nonlinear dimensionality reduction technique that finds a low-dimensional representation of graph structures [13, 44]. Though closely related to other manifold learning techniques including e.g., Laplacian-based spectral embedding, diffusion map embedding typically uses a different normalization process that approximates a Fokker-Planck diffusion equation instead of traditional laplacian normalization [59]. Moreover, diffusion map embedding attempts to model a multi-scale view of the diffusion process via a diffusion time parameter, t, that allow for more nuanced investigations of the geometry of the input data than are achievable via comparable techniques like spectral embedding. In the current manuscript we used diffusion time t = 0, which reveals the most global relationships of the input dataset [13].

Prior to embedding we thresholded our fused graphs, removing edges below the 90th percentile of weights for each individual, and computed the cosine similarity of the resulting network [54]. This network was decomposed using `mapalign` (https://github.com/satra/mapalign) to generate an embedded space for each of the 10,000 patient networks.

For our analyses we only considered embeddings for those networks in regions of parameter space deemed “stable” via the z-Rand thresholding procedure described in *Consensus clustering*, resulting in 1,262 embeddings. These embeddings were aligned using a generalized orthogonal Procrustes analysis (rotations and reflections only) and then averaged to generate a single patient embedding which was carried forward to all analyses (Fig. 1d). We only considered the first N components yielding a cumulative variance explained of ≥ 10%.

#### Embedding cross-validation

To examine the out-of-sample predictive utility of the dimensional embedding framework we re-ran the SNF grid search excluding two variables: the tremor dominant score (tremor) and the postural instability/gait difficulty score (PIGD). Embedded dimensions were regenerated following the same procedures depicted in Fig. 1d. We used a five-fold cross-validation framework to assess the extent to which embeddings predicted patient scores on the held-out clinical variables. A simple linear regression was performed using the first five embedding dimensions as predictors of each clinical score; betas were estimated from 80% of the patients and applied to predict scores for the remaining 20% of the patients. Predicted scores were correlated with actual scores, and the average correlations across all folds were reported.

### Statistical assessments

Quantitative assessments of the similarity between clustering assignments were calculated using normalized mutual information (NMI; [77]), a measure ranging from 0 to 1 where higher values indicate increased overlap between assignments. Alternative measures (e.g., adjusted mutual information) that correct for differing cluster numbers were not used as NMI was only calculated between assignments with the same number of clusters. Modularity was used to measure the “goodness-of-fit” of clustering assignments, and was calculated by:

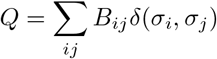

where *δ*() is the Kronecker delta function which returns 1 when patients *i* and *j* belong to the same cluster and 0 otherwise. Here, *B_ij_* is a modularity matrix whose elements are given by *B_ij_* = *A_ij_ − P_ij_*, where *A_ij_* and *P_ij_* are the observed and expected similarity between patients *i* and *j*. Modularity ranges between −1 and 1, where higher values indicate increasingly well-defined assortative clustering assignments [61].

To assess whether identified clusters were discriminable we performed separate one-way ANOVAs across groups for all data features provided to SNF using scipy’s statistical computing module [86] (model: `score ~ biotype`). Reported results were FDR-corrected (*q <* 0.05) using the Benjamini-Hochberg procedure [7] from the `statsmodels` modules [72]. Although longitudinal data were limited for many features, clinical-behavioral assessments were available for the majority of subjects up to five years post-baseline. Thus, we analyzed the impact of cluster affiliation on longitudinal feature scores using linear mixed effects models, including age, education, and sex as additional covariates.

## AUTHOR CONTRIBUTIONS

RDM and BM conceived the project. RDM designed and performed all analyses, with input from GS and CT. RDM and BM wrote the manuscript. AD and RBP provided input on all analyses and edited the manuscript.

## COMPETING INTERESTS

RBP personal fees from Takeda, Roche/Prothena, Novartis Canada, Biogen, Theranexus, GE HealthCare, Abbvie, and Otsuko, outside the submitted work.

## ACKNOWLEDGEMENTS

This research was undertaken thanks in part to funding from the Canada First Research Excellence Fund, awarded to McGill University for the Healthy Brains for Healthy Lives initiative. BM acknowledges support from the Canadian Insititutes of Health Research (CIHR-PJT) and from the Fonds de recherche du Québec Santé (Chercheur Boursier). RDM acknowledges support from the Healthy Brains for Healthy Lives (HBHL) initiative at McGill University and the Fonds de recherche du Québec Nature et technologies (FRQNT).

Data used in the preparation of this article were obtained from the Parkinson’s Progression Markers Initiative (PPMI) database (https://www.ppmi-info.org/data). For up-to-date information on the study, visit https://www.ppmi-info.org. The PPMI—a public-private partnership—is funded by the Michael J. Fox Foundation for Parkinson’s Research and funding partners, including AbbVie, Avid Radiopharmaceuticals, Biogen, BioLegend, Bristol-Myers Squibb, GE Healthcare, Genentech, GlaxoSmithKline (GSK), Eli Lilly and Company, Lundbeck, Merck, Meso Scale Discovery (MSD), Pfizer, Piramal Imaging, Roche, Sanofi Genzyme, Servier, Takeda, Teva, and UCB (https://www.ppmi-info.org/fundingpartners).

## SUPPLEMENTARY MATERIALS AND RESULTS

### SNF discriminates diagnostic groups

While the current report was primarily interested in investigating how SNF might serve to uncover putative biotypes of PD, we wanted to validate the use of SNF on identification of known diagnostic clusters in the same dataset. We ran the full SNF pipeline as described in the main text (see Fig. 1) including both healthy controls (*n* = 87) and PD patients (*n* = 186), to determine if SNF was able to correctly dissociate these groups. The resulting consensus cluster assignments yielded three clusters of *n* = 98, 84, and 91 individuals (Fig. S1); the third cluster (*n* = 91 individuals) showed strong overlap with the healthy control population (F1 score = 0.94), while the other two clusters split the PD population (combining the two clusters yielded an F1 score of 0.97). Notably, the healthy control cluster is easily distinguishable from the PD patient clusters along the first dimension of the embedding space (Fig. S1). This is quite distinct from the seemingly arbitrary groupings the PD biotypes make in the embedded space shown in Fig. 4, supporting the notion that biotypes may not be the most parsimonious characterization of PD (see *Discussion: Biotypes or dimensions?*).

### Assessing longitudinal clinical outcomes

To better investigate the clinical utility of the biotypes described in the main text we compared longitudinal clinical outcomes for these biotypes with those derived from different subsets of the data and other clustering techniques. Though there are numerous clinical variables to examine, we chose to focus on the two highlighted in the main text: the tremor dominant score (tremor) and the postural instability/gait difficulty score (PIGD).

First, we examined whether biotypes obtained solely from baseline clinical assessments—excluding CSF assays, DAT scans, and neuroimaging data—showed differences in clinical outcomes over time for these two variables. Clustering labels were derived using the same SNF grid search pipeline described in *Results: Similarity network fusion provides a viable alternative to data concatenation*, but using only baseline clinical assessments as input. We observe trends suggesting that PIGD scores are discriminable between biotypes at baseline, and there are no changes in this discriminability over time (Fig. S3b); tremor scores appear to be largely overlapping between biotypes.

Next, we examined whether biotypes derived from clustering using concatenated data showed discriminable clinical outcomes. These biotypes showed limited differences in PIGD and tremor scores at baseline, and relatively limited changes in discriminability over time (Fig. S3c). Interpretation of these results are supported by model estimates from linear mixed effects models run for both biotype definitions (Table S5).

### PCA is biased by data dimensionality

In the current article we compared clustering of PD patient data using SNF and data concatenation, showing that SNF better integrates data from different modalities and is less influenced by input data dimensionality. Here, we examine a similar comparison between the diffusion map embedding results presented in the main text and another low-dimensional projection technique: principal components analysis (PCA).

First, we computed the PCA of the concatenated patient by feature data matrix, yielding a vector of eigenvalues and corresponding eigenvectors. To investigate the similarities between this projection and the diffusion map embedding results we plotted the patient scores projected onto the first two principal components (PCs; Fig. S2a). Examining these projections when colored by patient affiliation with the PD biotypes reported in the main text reveals a much less distinct separation between biotypes than is observed with the diffusion map embedding projection (Fig. 4b). We confirmed these differences between projections by correlating patient scores along the first two PCs with their complementary scores along the first two dimensions of the diffusion map embedding, yielding low—but non-zero—correlations (*r*_PC1_ = 0.142, *r*_PC1_ = 0.266; Fig. S2b).

To investigate whether the PCA results were biased by data dimensionality we computed a PCA independently for each data modality and correlated patient scores along the first PC of these decompositions with the patient scores along the first PC of the concatenated data matrix (Fig. S2c). Notably, PC scores from the concatenated data matrix are almost perfectly correlated with PC scores along the first dimension of the cortical thickness data (*r* = 0.99998), while showing negligible correlation with PC scores from the other data modalities (*r*_avg_ = 0.01673). These results suggest that, like clustering, PCA of the concatenated data matrix is biased by data dimensionality and unsuitable for investigations of datasets like those presented in the current study.

### Stability of SNF

Despite extensive validation in the original article [87], its successful application to similar neuroimaging datasets [35, 76], and the exhaustive parameter search employed in the current article, we wanted to assess the stability of SNF as it applies to the current dataset. We first compare clustering assignments generated from SNF when varying the dimensionality of cortical thickness data (see *Data dimensionality variation*). We then compare patient networks generated from four hyperparameter combinations, examining differences in the resulting clustering solutions and diffusion dimensions (see *Hyperparameter variation*). Finally, we investigate how our choice of distance metric (i.e., squared Euclidean distance) impacts the patient networks and results reported in the main text (see *Alternative distance metrics*).

For all analyses, we calculated normalized mutual information scores (NMI) between different clustering solutions, and used Pearson correlations to compare the diffusion embedding dimensions derived from the fused netowkrs.

#### Hyperparameter variation

Using the results of the parameter search described in *Network creation and fusion* we selected four “stable” regions of parameter space and investigated the extent to which resulting clusters and embedded representations varied across these regions. For each of the four selected supra-threshold regions we calculated the center-of-mass and extracted the relevant (1) cluster labels and (2) network embeddings for the specified hyperparameter combinations (*K* = 22, 44, 58, 88; *μ* = 2.6, 4.4, 6.3, 7.3).

We computed NMI between all pairs of hyperparameter combinations separately for each cluster number (*n* = 2, 3, 4), finding considerable consistency amongst the three-cluster solutions (NMI_*three*_ = 0.71 0.08 [0.63– 0.86]) but notably lower consistency for the other cluster numbers (NMI_*two*_ = 0.49 0.15 [0.27–0.66]; NMI_*four*_ = 0.55 0.32 [0.26–0.92]). Embedding dimensions were very consistent across all parameter combinations (average *r* = 0.97 ± 0.03 [0.91–1.00]).

#### Alternative distance metrics

In the main text we use a squared Euclidean distance function to generate patient similarity networks from the original data feature matrices. While there is precedent for this choice [35, 76, 87], we also wanted to investigate the impact of the chosen distance metric on our results. Thus, we repeated the analyses described in the main text using two alternative distance metrics— cityblock (or Manhattan) distance and cosine distance– finding that both clustering results (NMI_*cityblock*_ = 0.74, NMI_*cosine*_ = 0.68) and embedding dimensions (average *r_cityblock_* = 0.93 0.10 [0.65–0.99], *r_cosine_* = 0.87s 0.10 [0.70–0.96]) were highly consistent with squared Euclidean distance.

### Comparing clustering techniques

Clinical clustering (“subtyping”) of individuals with Parkinson’s disease has received significant attention in recent years, especially since the release of the PPMI dataset [22, 24]. In the main text we demonstrated that inclusion of neuroimaging data significantly contributed to biotype discriminability using similarity network fusion. Here, we compare these findings to the clustering framework of one recent clustering solution using the same dataset.

#### Clinical guidelines (Fereshtehnejad et al., 2017)

In 2017, Fereshtehnejad and colleagues [24] used hierarchical clustering of baseline clinical-behavioral assessments to find three subtypes of individuals with PD from the PPMI dataset. From this clustering procedure they generated a set of clinical guidelines for classifying patients into three groups. While the hierarchical clustering and clinical guidelines result in slightly different subgroup definitions, the authors contend that their guidelines are more robust to different data acquisition schemes. Indeed, these guidelines have already been successfully applied to other datasets [17] with replication of primary clinical group differences. Although the replications studies report limited differences on autopsy parameters between groups, they attribute this to a ceiling effect driven by the advanced stage of the disease at death.

We apply these clinical guidelines to cluster our patient sample and find a similar distribution of assignments to those reported in [24] (90 subjects classified as “mild motor-predominant”, 68 as “intermediate”, and 28 as “diffuse malignant”). Notably, these assignments have limited overlap with the SNF-derived clustering assignments reported in the main text (normalized mutual information = 0.026). Using univariate one-way ANOVAs we find patients grouped according to the criteria of Fereshtehnejad are only discriminable in the clinical-behavioral assessments on which the clusters were initially defined (seven features in total; FDR-corrected, *q* < 0.05). That is, we fail to observe any cluster differences in DAT binding, CSF assays, cortical thickness, or subcortical volume measurements; we also fail to observe cluster differences in the PD-ICA atrophy scores derived from [97] (see *Materials and Methods: PD-ICA Atrophy Calculation*).

---

**Figure S1.**
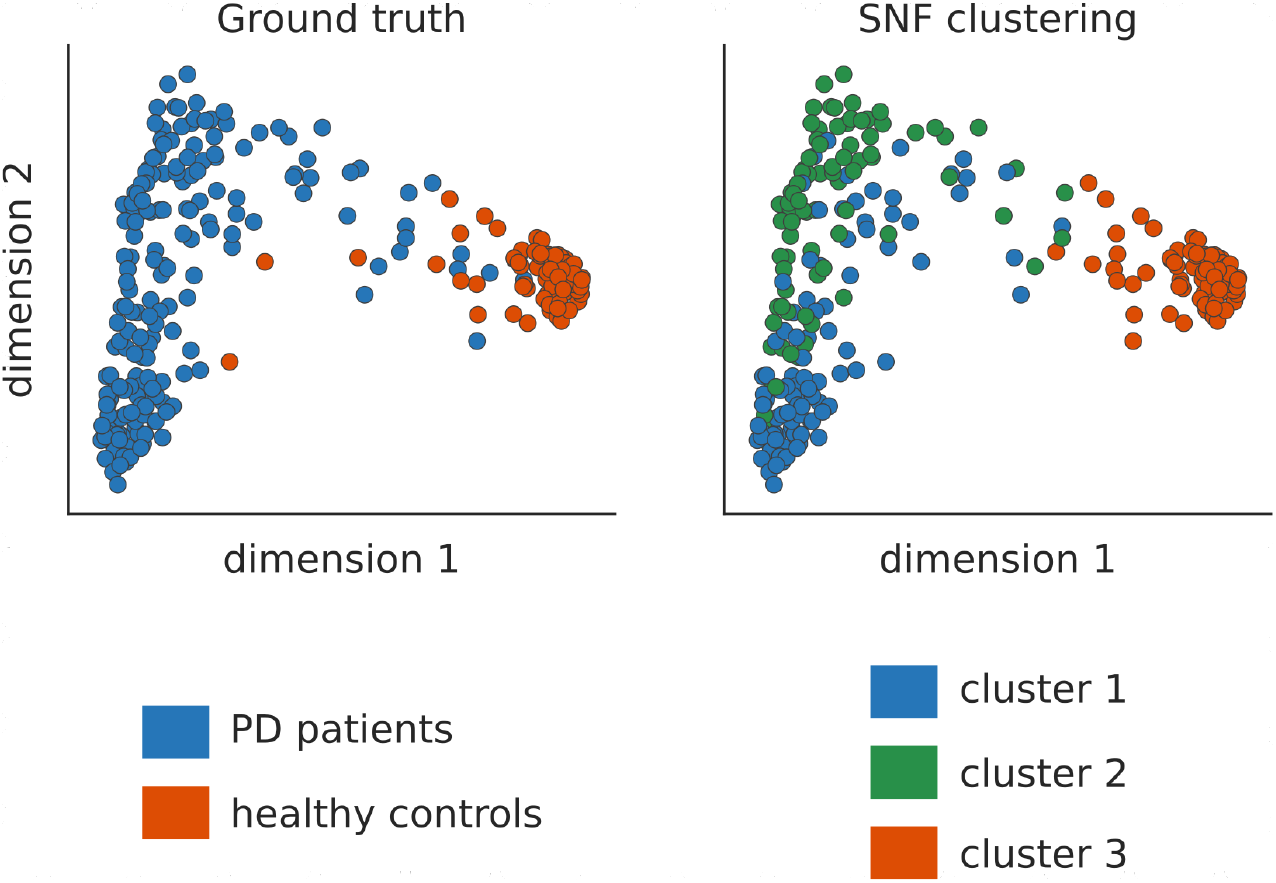
Similarity network fusion discriminates PD patients and healthy controls. True diagnostic groups (left panel; PD patients and healthy controls) compared with SNF-derived clustering assignments (right panel), plotted along the first two dimensions of the embedding space derived from SNF. Healthy individuals are largely assigned to their own cluster by SNF (cluster 3, orange), whereas PD patients are split amongst two clusters (cluster 1, blue, and cluster 2, green).

**Figure S2.**
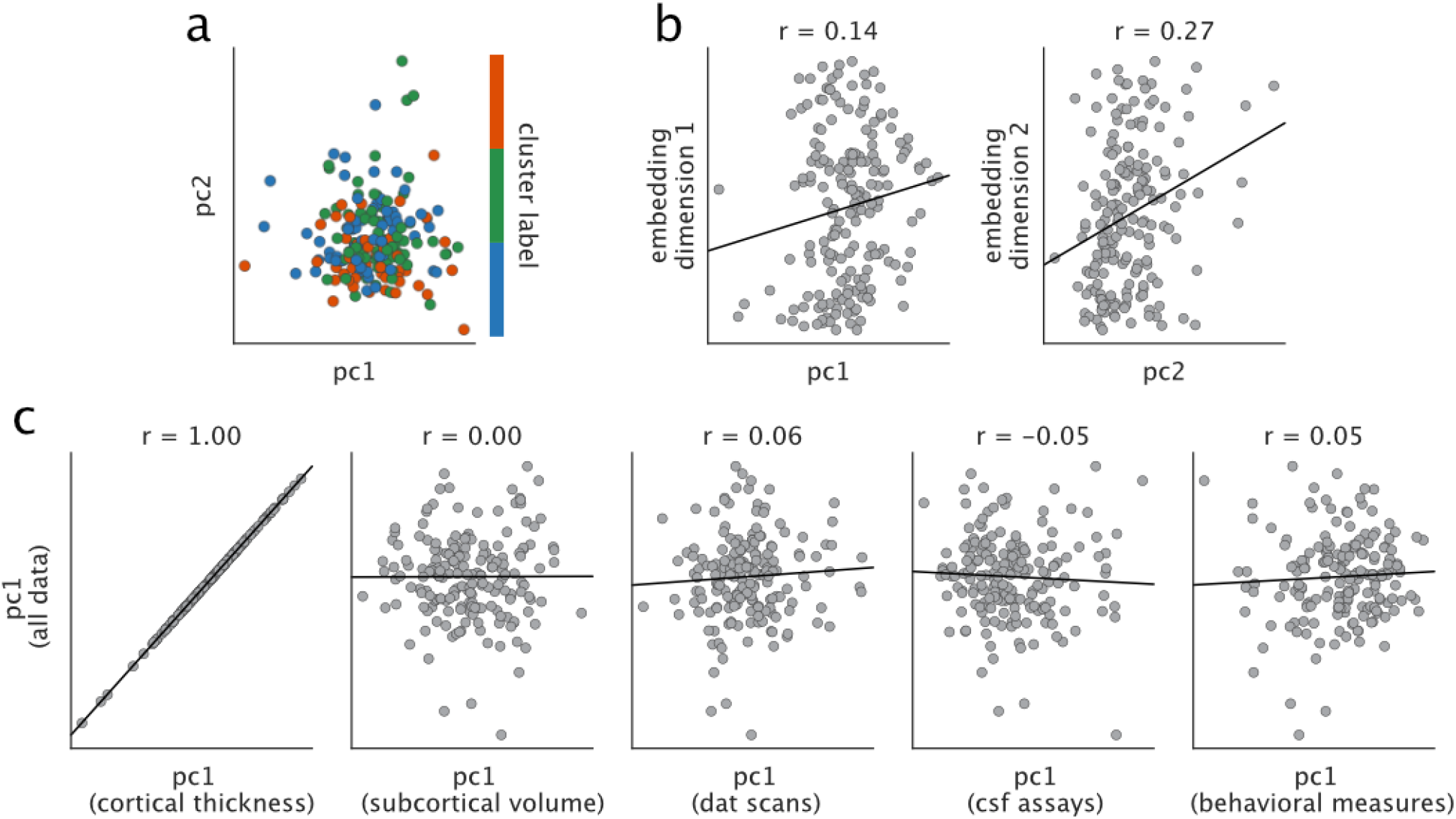
Comparison of low-dimensional embedding for SNF and data concatenation. **(a)** Projection of PD patients onto the first two principal components derived from the concatenated data matrix. Patients are colored by their affiliation to the biotypes presented in the main text. **(b)** Correlations of patient scores along the first two principal components with corresponding scores along the first two diffusion map embedding dimensions. **(c)** Correlations of patient scores along the first principal component from the concatenated data matrix with corresponding scores along the first principal component of each independent data modality.

**Figure S3.**
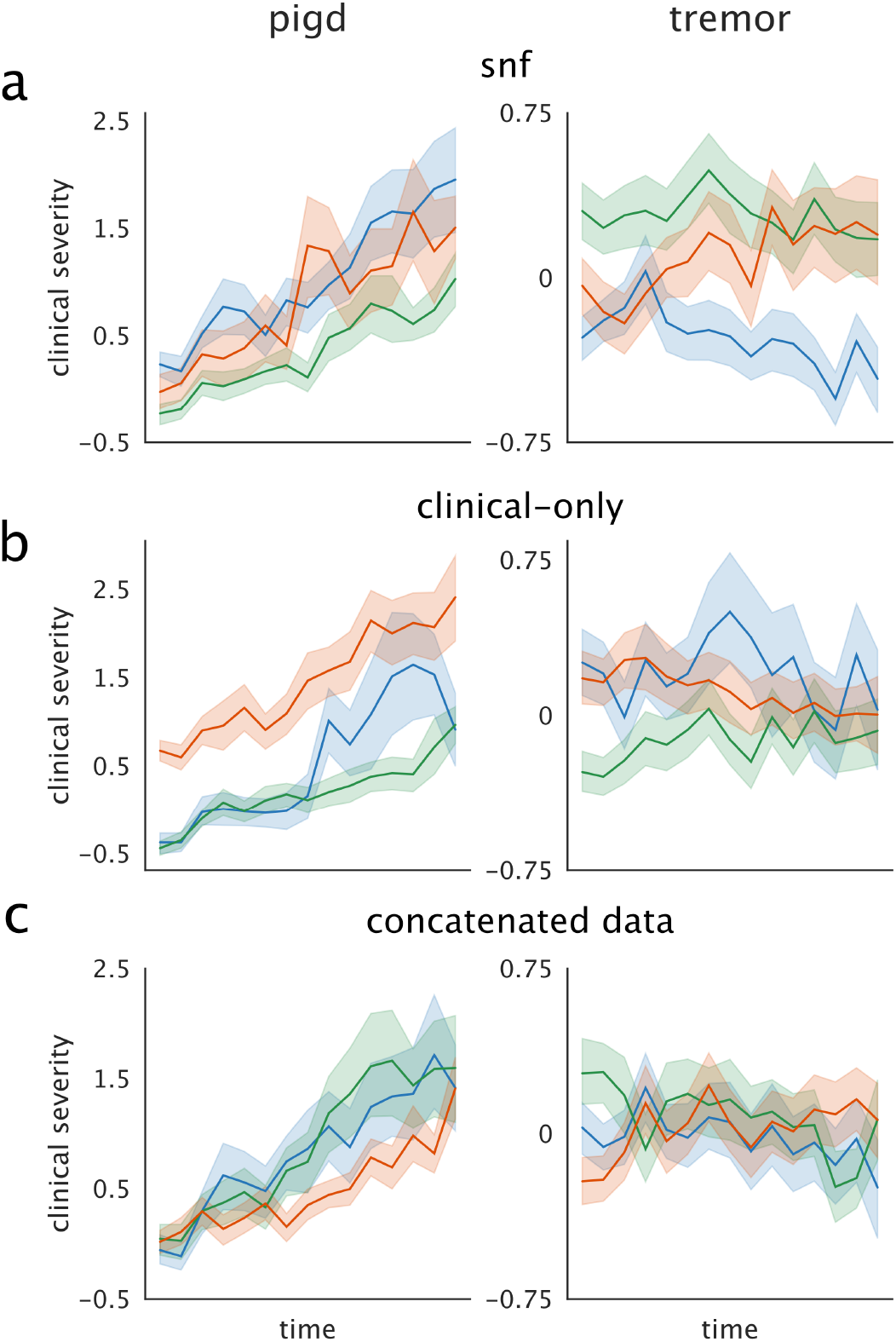
Longitudinal clinical outcomes vary based on data and clustering method. **(a)** Reproduction of panel (e) from Fig. 3 showing longitudinal trajectories of SNF-derived biotypes for postural instability/gait disorder (PIGD) and tremor scores. **(b)** Longitudinal trajectories for biotypes derived from baseline clinical assessments only. **(c)** Longitudinal trajectories for biotypes derived from concatenated data.

**TABLE S1.** Data features supplied to SNF. Cortical thickness features are not listed here due to their size, but can be found in the supplementary data included with this manuscript. Designations of (+) and (–) indicate approximate phenotypic severity, where (+) denotes that higher scores for the relevant feature are considered “more severe” and (–) denotes that lower scores for the relevant feature are considered “more severe”. A designation of (na) indicates that the relevant feature does not necessarily have a direct relationship to phenotypic severity. For all cortical thickness, subcortical volume, and DAT binding features lower scores are considered “more severe”.

| Subcortical volume (–) | DAT binding (–) | CSF assays | Clinical assessments |
| --- | --- | --- | --- |
| Caudate nucleus | Left caudate | Aβ (1-42) (–) | Benton (–) |
| Extended amygdala | Right caudate | CSF α-synuclein (–) | Epworth (+) |
| Globus pallidus, external | Left putamen | mtDNA deletion (+) | GDS (+) |
| Globus pallidus, internal | Right putamen | MT-ND1 copy number (+) | HVLT recall (–) |
| Habenular nuclei |  | MT-ND4 copy number (+) | HVLT recognition (–) |
| Hypothalamus |  | Phosphorylated tau (–) | HVLT retention (–) |
| Mammillary nucleus |  | Serum IGF-1 (–) | LNS (–) |
| Nucleus accumbens |  | Total tau (–) | MoCA (–) |
| Parabrachial pigmented nucleus |  |  | PIGD (+) |
| Putamen |  |  | QUIP (+) |
| Red nucleus |  |  | RBD (+) |
| Substantia nigra, pars compacta |  |  | SCOPA-AUT (+) |
| Substantia nigra, pars reticulata |  |  | Semantic fluency (–) |
| Subthalamic nucleus |  |  | STAI state (na) |
| Ventral pallidum |  |  | STAI trait (na) |
| Ventral tegmental area |  |  | Symbol digit (–) |
|  |  |  | Systolic BP drop (+) |
|  |  |  | Tremor (+) |
|  |  |  | UPDRS-I (+) |
|  |  |  | UPDRS-II (+) |
|  |  |  | UPDRS-III (+) |
|  |  |  | UPSIT (–) |

**TABLE S2.** Cluster demographics. Demographic information on clusters defined in *Results: Derived patient biotypes are clinically discriminable across modalities*. Presented values are means *±* standard deviations unless otherwise noted.

| Cluster | Age (yrs) | Male (%) | White (%) | Education (yrs) |
| --- | --- | --- | --- | --- |
| 1 | 60.24 ± 8.62 | 51.39 | 91.67 | 15.47 ± 2.78 |
| 2 | 62.29 ± 10.17 | 60.87 | 92.75 | 15.81 ± 2.81 |
| 3 | 62.69 ± 8.83 | 64.44 | 88.89 | 15.56 ± 3.11 |

| Cluster | Symptom duration (mo) | Family history of PD (%) | UPDRS total |
| --- | --- | --- | --- |
| 1 | 6.21 ± 6.41 | 18.06 | 34.04 ± 13.35 |
| 2 | 5.74 ± 5.45 | 31.88 | 31.09 ± 13.37 |
| 3 | 5.56 ± 6.28 | 22.22 | 29.93 ± 11.66 |

**TABLE S3.** Data features discriminable between SNF-derived patient clusters. Only features significant after FDR-correction (*q* < 0.05) are reported; note that no clinical-behavioral assessments survived this threshold. Numbers in parentheses next to cortical thickness features indicate regional sub-divisions; for more information on the parcellation refer to [11].

| Cortical thickness | Subcortical volume | DAT binding | CSF assays |
| --- | --- | --- | --- |
| Right supramarginal gyrus (15) | Substantia nigra, pars compacta | Left caudate | Total tau |
| Right superior frontal gyrus (13) | Red nucleus | Left putamen | Phosphorylated tau |
| Right lateral orbitofrontal cortex (14) | Subthalamic nucleus | Right caudate | CSF α-synuclein |
| Right isthmus cingulate (5) | Parabrachial pigmented nucleus | Right putamen |  |
| Right medial orbitofrontal cortex (10) | Ventral pallidum |  |  |
| Right insular cortex (14) | Globus pallidus, internal |  |  |
| Right superior frontal gyrus (20) | Substantia nigra, pars reticulata |  |  |
| Left paracentral lobule (10) | Hypothalamus |  |  |
| Left superior temporal gyrus (25) | Extended amygdala |  |  |
| Left insular cortex (6) | Globus pallidus, external |  |  |
|  | Ventral tegmental area |  |  |
|  | Nucleus accumbens |  |  |
|  | Putamen |  |  |
|  | Habenular nuclei |  |  |

**TABLE S4.** Linear mixed effect model of tremor and PIGD scores. Parameter estimates for two linear mixed effect models fit to longitudinal patient data with formula: `score ~ time * biotype + age + education + sex` [72]. Relevant data are shown in Figure 3.

| Tremor model |  |  |  |  |  |
| --- | --- | --- | --- | --- | --- |
|  | Estimate | SE | Z-stat | 2.5% CI | 97.5% CI |
| Intercept | -1.212 | 1.000 | -1.213 | -3.171 | 0.747 |
| Biotype [T.2] | 0.792 | 0.276 | 2.866 | 0.251 | 1.334 |
| Biotype [T.3] | 0.086 | 0.313 | 0.276 | -0.526 | 0.699 |
| Sex [M-F] | 0.170 | 0.238 | 0.717 | -0.295 | 0.636 |
| Time | -0.081 | 0.024 | -3.391 | -0.128 | -0.034 |
| Time:Biotype [T.2] | 0.074 | 0.034 | 2.193 | 0.008 | 0.140 |
| Time:Biotype [T.3] | 0.177 | 0.039 | 4.504 | 0.100 | 0.253 |
| Age | 0.042 | 0.013 | 3.378 | 0.018 | 0.067 |
| Education | 0.064 | 0.041 | 1.580 | -0.015 | 0.144 |

| PIGD model |  |  |  |  |  |
| --- | --- | --- | --- | --- | --- |
|  | Estimate | SE | Z-stat | 2.5% CI | 97.5% CI |
| Intercept | -0.581 | 0.474 | -1.225 | -1.509 | 0.348 |
| Biotype [T.2] | -0.281 | 0.132 | -2.127 | -0.540 | -0.022 |
| Biotype [T.3] | -0.147 | 0.150 | -0.984 | -0.440 | 0.146 |
| Sex [M-F] | -0.067 | 0.113 | -0.591 | -0.287 | 0.154 |
| Time | 0.166 | 0.013 | 12.937 | 0.141 | 0.191 |
| Time:Biotype [T.2] | -0.046 | 0.018 | -2.533 | -0.081 | -0.010 |
| Time:Biotype [T.3] | 0.017 | 0.021 | 0.827 | -0.024 | 0.058 |
| Age | 0.023 | 0.006 | 3.879 | 0.011 | 0.035 |
| Education | -0.006 | 0.019 | 0.770 | -0.043 | 0.032 |

**TABLE S5.**
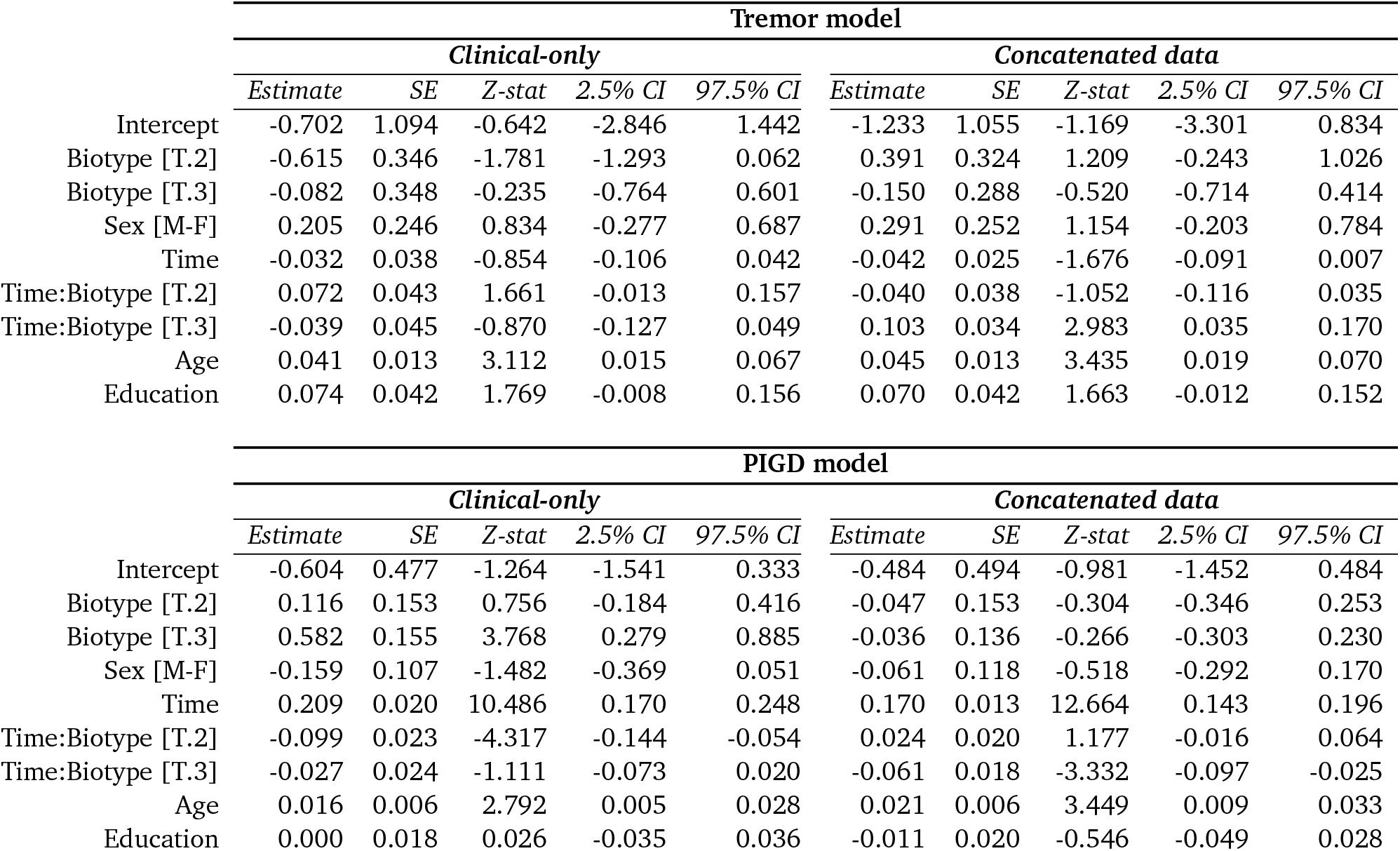
Longitudinal clinical outcomes vary based on data and clustering method. Parameter estimates for supplementary linear mixed effect models fit to longitudinal patient data with formula: `score ~ time * biotype + age + education + sex` [72]. Biotypes are derived from either baseline clinical data only (“clinical-only”) or concatenated multimodal data (“concatenated data”). Relevant data are shown in Figure S3b-c.

| Tremor model |  |  |  |  |  |  |  |  |  |  |
| --- | --- | --- | --- | --- | --- | --- | --- | --- | --- | --- |
|  | Clinical-only |  |  |  |  | Concatenated data |  |  |  |  |
|  | Estimate | SE | Z-stat | 2.5% CI | 97.5% CI | Estimate | SE | Z-stat | 2.5% CI | 97.5% CI |
| Intercept | -0.702 | 1.094 | -0.642 | -2.846 | 1.442 | -1.233 | 1.055 | -1.169 | -3.301 | 0.834 |
| Biotype [T.2] | -0.615 | 0.346 | -1.781 | -1.293 | 0.062 | 0.391 | 0.324 | 1.209 | -0.243 | 1.026 |
| Biotype [T.3] | -0.082 | 0.348 | -0.235 | -0.764 | 0.601 | -0.150 | 0.288 | -0.520 | -0.714 | 0.414 |
| Sex [M-F] | 0.205 | 0.246 | 0.834 | -0.277 | 0.687 | 0.291 | 0.252 | 1.154 | -0.203 | 0.784 |
| Time | -0.032 | 0.038 | -0.854 | -0.106 | 0.042 | -0.042 | 0.025 | -1.676 | -0.091 | 0.007 |
| Time:Biotype [T.2] | 0.072 | 0.043 | 1.661 | -0.013 | 0.157 | -0.040 | 0.038 | -1.052 | -0.116 | 0.035 |
| Time:Biotype [T.3] | -0.039 | 0.045 | -0.870 | -0.127 | 0.049 | 0.103 | 0.034 | 2.983 | 0.035 | 0.170 |
| Age | 0.041 | 0.013 | 3.112 | 0.015 | 0.067 | 0.045 | 0.013 | 3.435 | 0.019 | 0.070 |
| Education | 0.074 | 0.042 | 1.769 | -0.008 | 0.156 | 0.070 | 0.042 | 1.663 | -0.012 | 0.152 |

| PIGD model |  |  |  |  |  |  |  |  |  |  |
| --- | --- | --- | --- | --- | --- | --- | --- | --- | --- | --- |
|  | Clinical-only |  |  |  |  | Concatenated data |  |  |  |  |
|  | Estimate | SE | Z-stat | 2.5% CI | 97.5% CI | Estimate | SE | Z-stat | 2.5% CI | 97.5% CI |
| Intercept | -0.604 | 0.477 | -1.264 | -1.541 | 0.333 | -0.484 | 0.494 | -0.981 | -1.452 | 0.484 |
| Biotype [T.2] | 0.116 | 0.153 | 0.756 | -0.184 | 0.416 | -0.047 | 0.153 | -0.304 | -0.346 | 0.253 |
| Biotype [T.3] | 0.582 | 0.155 | 3.768 | 0.279 | 0.885 | -0.036 | 0.136 | -0.266 | -0.303 | 0.230 |
| Sex [M-F] | -0.159 | 0.107 | -1.482 | -0.369 | 0.051 | -0.061 | 0.118 | -0.518 | -0.292 | 0.170 |
| Time | 0.209 | 0.020 | 10.486 | 0.170 | 0.248 | 0.170 | 0.013 | 12.664 | 0.143 | 0.196 |
| Time:Biotype [T.2] | -0.099 | 0.023 | -4.317 | -0.144 | -0.054 | 0.024 | 0.020 | 1.177 | -0.016 | 0.064 |
| Time:Biotype [T.3] | -0.027 | 0.024 | -1.111 | -0.073 | 0.020 | -0.061 | 0.018 | -3.332 | -0.097 | -0.025 |
| Age | 0.016 | 0.006 | 2.792 | 0.005 | 0.028 | 0.021 | 0.006 | 3.449 | 0.009 | 0.033 |
| Education | 0.000 | 0.018 | 0.026 | -0.035 | 0.036 | -0.011 | 0.020 | -0.546 | -0.049 | 0.028 |

## Notes

https://github.com/netneurolab/markello_ppmisnf

### Competing Interest Statement

The authors have declared no competing interest.

